# High-throughput, single-cell-encoded colony imaging reveals infection-linked heterogeneity

**DOI:** 10.64898/2026.11.03.764039

**Authors:** Sena Gizem Suer, Tugrul Doruk, Narmin Hasanzada, Karl Kochanowski, Kemal Avican

## Abstract

Phenotypic heterogeneity within bacterial populations shapes adaptation, antibiotic tolerance, and infection dynamics, yet colony-level quantification remains difficult at scale and on dark/selective media. We present HeteroTyper, an automated, high-throughput time-lapse platform that measures colony phenotypes across >100 plates per run on both transparent and dark media. HeteroTyper includes a ready-to-use computational pipeline that can be deployed on common plate-imaging setups and across agar types, delivering appearance times as a proxy for lag times, growth trajectories, colony size, intensity, and morphology from routine image series. Automated counts closely matched manual counts, supporting accuracy and scalability. Using *Yersinia pseudotuberculosis* as a model, HeteroTyper resolves phenotypic heterogeneity *in vitro*, with early and late phases showing broader appearance time distributions than mid-log phases. It also reveals organ-specific heterogeneity *in vivo*, with mesenteric lymph node isolates displaying the broadest appearance time distributions compared to spleen and liver. Appearance time was negatively associated with final colony size and intensity, linking growth initiation to downstream colony outcomes. Together, these distribution-level measurements indicate that bacterial adaptation during infection is compartmentalized by tissue microenvironments and prior history, and establish HeteroTyper as a practical, generalizable route to quantify infection-relevant phenotypic diversity on solid media at routine, high throughput.

**Importance:** Understanding why some bacteria in the same population grow quickly while others wait is key to tackling infections and treatment failure. However, most tools cannot measure these differences at scale or on the dark, selective agars used in clinical microbiology. HeteroTyper is a simple, automated platform that time-lapse images more than a hundred plates at once and quantifies when colonies first appear (appearance time), how fast they grow, and how they look. Using HeteroTyper, we show that bacterial behavior depends on prior growth history and on the organ they come from. Mesenteric lymph node isolates display the broadest spread of appearance times, whereas spleen and liver isolates are more uniform. By making realistic, high-throughput measurements routine, HeteroTyper enables laboratories to detect hard- to-see subpopulations that may underlie antibiotic tolerance and relapse.

## Introduction

Phenotypic heterogeneity, a widespread phenomenon in microbial populations, refers to a diverse range of traits exhibited by cells within a single population, particularly in pathogenic species. This variability enables pathogen subpopulations to adapt effectively to the complex and fluctuating environments encountered within a host^1^ and can be structured by tissue-specific microenvironments during infection, leading to co-existing slow- and fast-growing subpopulations^2,3^. Besides environmental factors, the heterogeneity may arise due to genetic mutations or epigenetic modifications^4,5^ that can contribute to the emergence of antibiotic-resistant subpopulations^6^. When exposed to antibiotics, these resistant subpopulations survive, leading to treatment failure and recurrent infections and, more broadly, antibiotic-tolerant persister states that complicate therapy^7^. Phenotypes can be quantitatively assessed by observing variations across a spectrum of parameters, including growth rate, lag time, antibiotic susceptibility, and stress response^4,5,8^. Among these, differences in lag times during growth for distinct bacterial subpopulations represent a crucial indicator of heterogeneity^9-11^ and can be adaptively tuned to enhance survival under intermittent antibiotic exposure^10-13^. Notably, prolonged bacterial lag times are associated with the emergence of small colony variants (SCVs), which constitute a subpopulation of persister cells that exhibit tolerance to antibiotic treatments despite lacking genetic resistance^14^. These SCVs can arise *de novo* under various stress conditions, including those found in abscesses or acidic media, and their prevalence has been correlated with antibiotic usage^9,10,15^. Therefore, identification of cells whose lag time varies among others within a population is essential for elucidating infection pathogenesis, identifying the mechanisms of antimicrobial resistance, and ultimately developing more effective therapeutic strategies targeting pathogens. However, lack of quantitative and scalable approaches to analyse phenotypically heterogeneous populations, especially within dense microbial communities or *in situ* during infection leaves a significant gap in microbiology.

While numerous methodologies have been established for analysing bacterial colony growth parameters, including lag time, their scalability and throughput often present limitations^10,11,16^. For instance, CellProfiler is capable of analysing colony features from agar plate images, but it is restricted to processing one experimental image per run^17^. This single-image analysis per run significantly impedes high-throughput investigations by prolonging processing times across multiple plates, highlighting the need for automated image analysis solutions that enable rapid processing. Similarly, other approaches, such as those employed to characterize bacterial lag times in *Listeria monocytogenes*, have been constrained to a limited number of plates per experiment^18^. On the other hand, techniques like ScanLag^11^ offer speed and scalability but rely on scanning from bottom, restricting their applications to transparent agar. In dark or opaque media, reduced light transmission disrupts image acquisition and precise detection of colony features, rendering them unsuitable for many clinically relevant pathogens requiring such complex, media types. Additional colony-analysis tools demonstrate the need for scalable, generalizable pipelines^19^, while current technics largely rely on transparent media^20^. Related automated colony quantification systems further highlight the long-standing demand for reliable image-to-phenotype workflows^21^. Camera-based methods effectively analyse heterogeneity in clinical isolates on diverse agar type^10,18^. However, their design, typically employing one camera per plate, limits throughput. Addressing the technical challenges of quantitatively assessing phenotypic heterogeneity at single-cell resolution, there is a clear demand for high-throughput methodologies capable of managing large numbers of agar plates and providing exhaustive insights into bacterial growth dynamics and heterogeneity.

We introduce HeteroTyper, a high-throughput system that overcomes the dependence on specific media types by acquiring images from above and enables high-throughput analysis of bacterial growth and heterogeneity across numerous agar plates. The HeteroTyper system utilizes a movable camera to image up to 104 plates in a single experiment and captures time lapse images to identify phenotypic heterogeneity in a bacterial population by measuring appearance times of tens of thousands of individual colonies. Imaging from above with a camera mounted on a guide rail enables fast image acquisition and helps distinguish true colonies from the matte-black background, thereby improving the performance of current segmentation pipelines which are implemented in HeteroTyper. In this study, we used colony appearance time as a proxy for bacterial lag times as it was proposed in multiple reports^10,11^. The universal MATLAB pipeline applies fixed preprocessing and segmentation parameters to extract colony features across diverse agar media.

In this study, *Yersinia pseudotuberculosis* was used as a model organism to investigate bacterial heterogeneity at single cell level during *in vitro* growth and colonization in different host organs including mesenteric lymph nodes (MLNs), spleen, and liver. We detected variations in appearance time, colony size, and intensity across both *in vitro* and *in vivo* populations at high throughput using either bright or dark agar plates. Appearance time distributions revealed more heterogeneous profiles for colonies originating from early or late growth phases, where early-appearing colonies often exhibited higher intensities. *In vivo* samples were also highlighted by diverse appearance time distributions and morphological appearances, indicating variabilities in bacterial adaptation to different infection environments. Overall, HeteroTyper provides an accurate, quantitative and high-throughput analysis of bacterial population heterogeneity, offering valuable insights into bacterial growth dynamics under different conditions.

## Results

### Design and automation of the HeteroTyper system

To enable high-throughput, time-resolved monitoring of bacterial heterogeneity at single cell level in a population, we developed HeteroTyper system. The system repurposes the motion-controlled gantry of a laser cutter, on which a GoPro Hero 10 camera is mounted, allowing precise X-Y traversal and overhead imaging of each plate **(Fig. 1A, Supplementary Figure S1)**. Imaging events are controlled by an Arduino Nano microcontroller, which regulates gantry movement and transmits a 5 V trigger signal via a remote control to initiate image capture. To facilitate colony segmentation, plates are positioned on a matte-black background, enhancing contrast between bacterial colonies and the background. Once initiated, the system proceeds autonomously, acquiring images at user-defined intervals over extended time periods. The imaging schedule is fully configurable through the Arduino interface, offering flexibility across a range of experimental conditions. Related high-throughput plate phenomics frameworks and transparent-agar time-lapse systems highlight the community need for scalable imaging, which HeteroTyper extends to dark/complex media^20,22^.

**Fig. 1.**
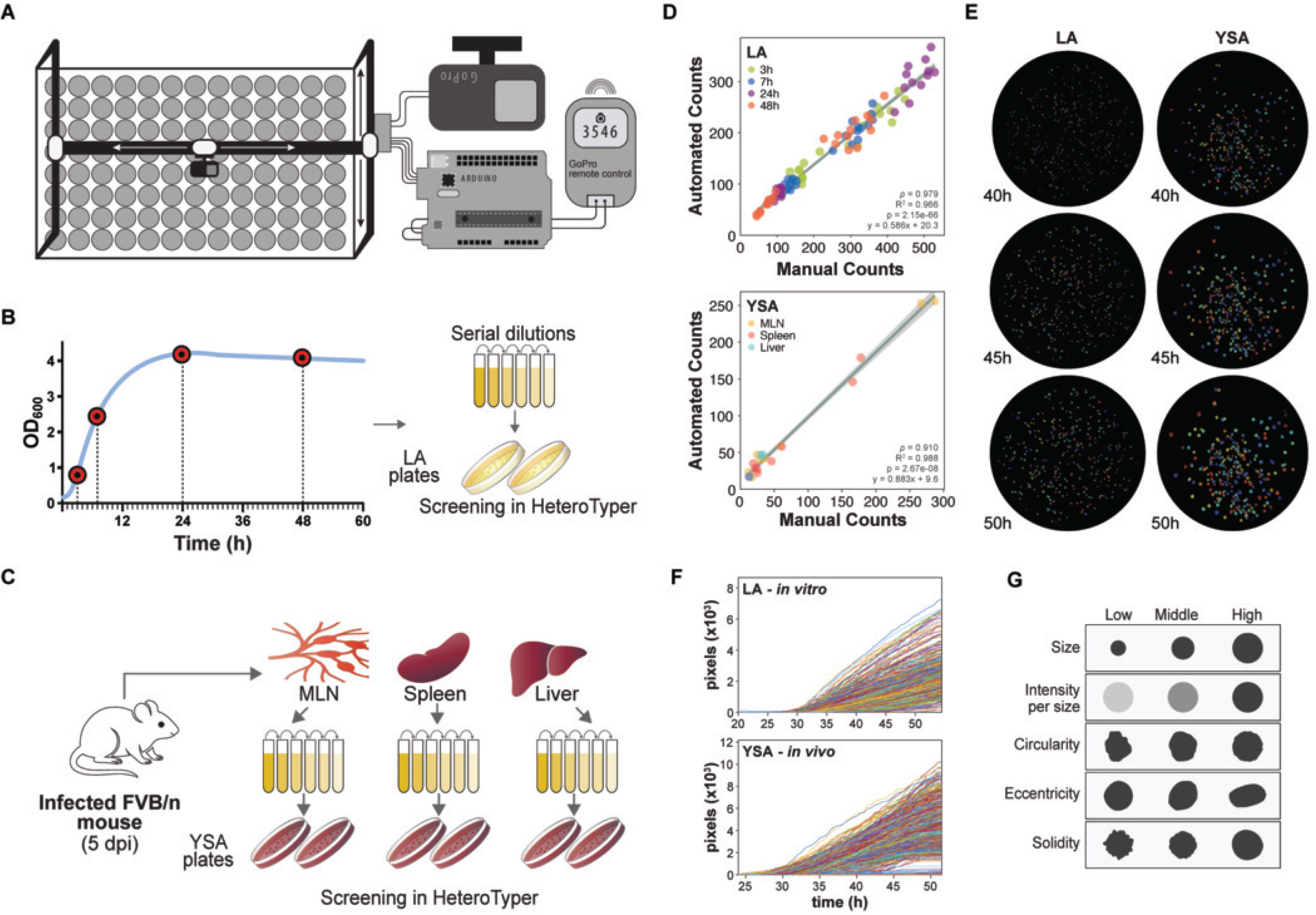
Design and experimental workflow of the HeteroTyper automated imaging system with capabilities. **A** View of HeteroTyper from above while the lid is open. The bed size of HeteroTyper is 102,5X152 cm allowing, 104 petri dishes to be screened. **B** Experimental setup for *in vitro Y. pseudotuberculosis* cells using freshly inoculated liquid culture from overnight in LB media. Samples were collected at different time points (3, 7, 24, and 48 hours) during growth, serially diluted, and plated on LA plates to obtain a range of colony numbers. These plates were then screened using the HeteroTyper system to monitor bacterial growth over time. **C** Experimental setup for *in vivo Y. pseudotuberculosis* cells. FVB\n mice were orally infected with *Y. pseudotuberculosis* YPIII, and mesenteric lymph nodes (MLN), spleen, and liver samples were collected at 5 dpi. The organs were homogenized, serially diluted, and plated on YSA plates before screening in the HeteroTyper system. **D** Comparison of manual and automated colony counts for LA and YSA plates demonstrating the accuracy of the computational analysis. Pearson correlation coefficients (r), coefficients of determination (R^2^), corresponding *p*-values (*p*), and fitted linear equations (y = mx + b) are displayed individually for each panel. **E** Segmented colony images from representative plates, showing *in vitro* (24 hours) colonies on LA and *in vivo* (MLN) colonies on YSA plates at different time points (40, 45, and 50 hours). **F** Growth curves of *in vitro* (24 hours) and *in vivo* (MLN) *Y. pseudotuberculosis* colonies were derived from all replicate plates of the respective sample groups. **G** Schematic illustration of colony morphologies showing low, middle, and high values for size, intensity per size, circularity, eccentricity, and solidity parameters.

The HeteroTyper computational pipeline extracts key phenotypic parameters from time-lapse images, including colony morphology, colony appearance time, size, and pixel intensity. These outputs enable the detection of heterogeneity in growth dynamics and appearance times among individual colonies. The analysis workflow begins with pre-processing steps that include detection of plate centers, image standardization, and colony segmentation. Briefly, images were pre-filtered to exclude plates with centers deviating >10% from the median position. After grayscale conversion, illumination was corrected using a Gaussian filter, followed by intensity normalization and contrast enhancement. Noise was reduced with Gaussian blur, and colonies were detected using a Laplacian of Gaussian–based threshold. Colonies were segmented by watershed and further filtered by size and shape to exclude noise, edge artifacts, and merged or non-circular objects (see methods). Pre-processing and segmentation choices were informed by established image-analysis pipelines widely used in microbiology^17,23,24^.

To benchmark the system, we used *Yersinia pseudotuberculosis* YPIII as a model organism and investigated heterogeneity in both *in vitro* cultures and bacteria recovered from infected mouse tissues. To assess the scalability of the system on both transparent and dark agar plates, we plated *in vitro* cultures on Luria Agar (LA) and tissue recovered bacteria to dark *Yersinia* Selective Agar (YSA). For *in vitro* analysis, we subcultured overnight *Y. pseudotuberculosis* in rich medium, sampled at 3, 7, 24, and 72 hours of growth, serially diluted, and plated onto eight LA plates per condition per biological replicate **(Fig. 1B)**. We performed *Y. pseudotuberculosis in vitro* growth experiments at 26°C as it exhibits growth restriction at 37°C. For the *in vivo* dataset, four FVB/n mice were orally infected with a low dose (10⁷ CFU/ml) of *Y. pseudotuberculosis*. At 5 days post-infection (dpi), MLNs, spleens, and livers were dissected, homogenized in PBS, serially diluted, and plated onto dark YSA **(Fig. 1C)**. Further analysis was performed on the resulting colonies. Manual and automated counts on both media were highly correlated (Spearman *ρ* ≥ 0.91; **Fig. 1D**), confirming reliable segmentation. To provide full transparency regarding absolute quantification, we fitted the regression lines, which had slopes of 0.586 for LA and 0.883 for YSA plates. These slope values indicate a modest, systematic undercount by the automated system. This difference is primarily due to the conservative exclusion of colonies located on the edges of the Petri dish during the segmentation process, an approach designed to prevent edge-effect artifacts and ensure the extraction of high-quality phenotypic data. Agreement between automated and manual colony quantification is consistent with prior benchmarks in open-source tools such as OpenCFU^19^. Time-lapse snapshots and single-colony trajectories **(Fig. 1E, F, Supplementary Figure S2A-D)** provide temporal quality control and reveal dynamic heterogeneity in lag, growth and morphology including colony size, intensity per size, circularity, eccentricity, and solidity **(Fig. 1G)**. We linked the distribution patterns to concrete phenotypes by showing schematic illustration of colonies **(Fig. 1G)** that represents low, middle, and high values for each colony morphology. Size represents the segmented colony area. Intensity is measured as the summed pixel intensity within the colony area, reflecting the optical density of the colony, that is, how much light is absorbed or scattered by its biomass. Since comparisons are made within the same species, differences in colony colour do not substantially influence this relationship. Intensity per size is then calculated by normalizing intensity to colony area, providing a measure of cell density per unit area, where higher values indicate thicker biomass accumulation. Circularity measures how closely a colony resembles a perfect circle, whereas eccentricity indicates how elongated a colony is. Solidity reflects how filled or compact the colony appears by comparing its actual area to the area it would have if its outline was perfectly smooth. Shape and texture descriptors similar to circularity, solidity and eccentricity are widely used in colony phenotyping pipelines^17,23-25^.

### HeteroTyper detects heterogeneity in populations by measuring appearance time

Variations in appearance time, reflected by delayed onset of growth, likely arise from physiological or genetic differences prior to plating, as well as microenvironmental effects. Such heterogeneity is indicative of functional diversity within the population. Appearance time was defined as the first time point at which colony area exceeded a user-defined threshold. Established systems such as ScanLag and ColTapp use pixel-based detection threshold, ScanLag typically defining colony appearance at a threshold of 10 pixels (px) and ColTapp reporting limitations for colonies with a radius below 3 px^11,26^. However, these thresholds depend on the imaging system and image quality. For HeteroTyper, we tested diverse colony size thresholds using the GoPro Hero 10 imaging setup and selected 100 px as the optimal threshold for reliable colony detection and appearance time determination **(Supplementary Figure S3)**. To assess the capabilities of the HeteroTyper in determining differences in heterogeneity within a growing bacterial population via appearance time distribution, we screened colonies obtained from WT *Y. pseudotuberculosis* culture at different growth phases and from different mouse organs on transparent LA and dark colored YSA plates, respectively. Colony appearance time as a plate-based proxy for lag time has been used in automated time-lapse systems^22^ and aligns with single-cell observations that lag time variability is biologically meaningful under stress and antibiotics^13^.

To accurately analyse *in vitro* samples, we screened dilution series ranging from 10^4^ to 10^7^ depending on time point and selected the plates with appropriate colony numbers (32-500 colonies). While the optimal count for *E. coli* is typically below 200 due to its larger colony size (2–4 mm at 37°C for 24 hours), the smaller diameter of *Y. pseudotuberculosis* (0.5–1.5 mm at room temperature for 48 hours) allows this higher range to remain adequate in 100-mm petri dishes used in the experiments. To quantify the variance within these populations, we utilized the Gini coefficient as a measure of phenotypic heterogeneity, as it provides a useful measure of inequality in a distribution regardless of absolute sample size or mean values. The results, incorporating phenotypic distributions and Gini coefficients, demonstrated that *Y. pseudotuberculosis* cells exhibit distinct profiles for specific features across different growth phases. Appearance time of colonies originating from 3- and 7-hour growth periods peaked earlier than those from 24- and 48-hour growth periods **(Fig. 2A)**. However, the appearance time distribution of 3- and 48-hour colonies exhibited wider ranges with longer tails (prolonged appearance time) supported by higher Gini coefficient, indicating greater heterogeneity at these time points **(Fig. 2A)**. To confirm that these Gini coefficients reflect the observed phenotypic heterogeneity and not driven by the number of colonies per plate, we examined the relationship between colony count and Gini coefficient for appearance time across all sample groups **(Supplementary Figure S4A)**. Spearman correlation revealed no significant association at any growth phases (*p* > 0.05), supporting that the heterogeneity captured in appearance time distributions is independent of the number of colonies identified per plate.

**Fig. 2.**
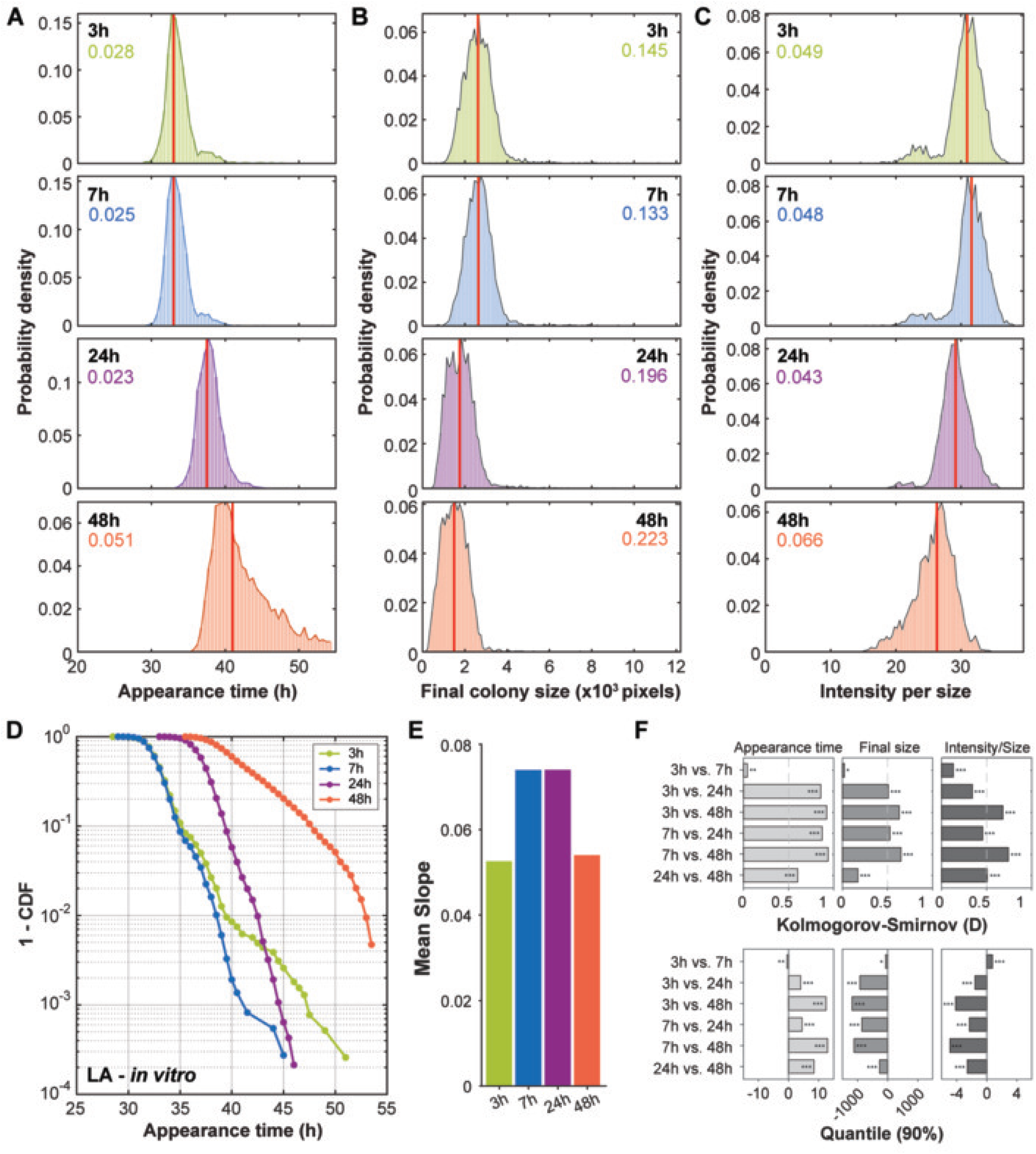
Variation in colony phenotypes of *Y. pseudotuberculosis* reflect levels of heterogeneity at different growth phases. For *in vitro* experiments, wt cells were inoculated into a rich liquid medium and grown at 26°C. Samples were collected from the same culture at 3, 7, 24, and 48 hours of growth, serially diluted and the 10^4-7^ dilutions were spread onto 8 plates for each time point analyses and replicate, incubated at room temperature for 20 hours, and imaged in the HeteroTyper system. **A** Appearance time, **B** final colony size, and **C** summed pixel intensity per colony size distributions using probability distribution function of bacterial colonies on agar plates originating from cells grown for 3h (n = 3869), 7h (n = 3654), 24h (n = 4593), and 48h (n = 3170). **D** Appearance times of *in vitro* (LA) cells are plotted as complementary cumulative distribution function (1–CDF) on logarithmic scale. **E** Mean slope values of each sample group represented in **D. F** Pairwise comparisons between growth conditions are represented for appearance time, final colony size, and intensity per size using bar plots. Kolmogorov–Smirnov statistics (D) are displayed in the upper panel, and differences in the 90th percentile are shown in the lower panel. Appearance time is represented in hours (h), with an additional incubation period of 20 hours, while the colony size was measured in pixels (px). For each parameter in **A-C**, Gini coefficients are shown in colors of growth condition and median values are indicated by red vertical lines. Asterisks indicate statistical significance (*p* <0.05: *, *p* < 0.01: **, *p* < 0.001: ***) in **F**.

Under conditions of starvation, such as bacterial cells at 48 hours of growth, appearance time prior to re-initiation of growth can be extended^11^. This prolonged appearance time occurred because the cells need additional time to recover and resume metabolic activities necessary for growth. Prolonged appearance time after nutrient limitation/starvation is consistent with prior observations linking growth history to delayed regrowth and tolerance^13^. In contrast, bacterial cultures at 3 hours generally had shorter appearance time and few colonies with long appearance times. This observation suggests that the 3-hour duration of growth is insufficient for a number of cells in the population to adapt and regrow in liquid medium when provided with fresh culture medium. These phenotypes were reflected with final colony size distribution at the end of 48 hours incubation on plates where populations with higher appearance times generally displayed lower median final colony sizes **(Fig. 2B)**. Interestingly, colonies originating from 7 hours growth were larger in size, indicating faster growth following shorter appearance time. To determine whether these differences reflect true statistical divergence rather than observable trends in median lines, we performed pairwise hypothesis testing across all sample groups for each phenotypic feature. Colony size and appearance time both differed significantly between the 7-hour and the 24- and 48-hour growth (Wilcoxon rank-sum, *p* < 1e-300 for both), supporting the association between shorter appearance time and larger final colony size **(Supplementary Table S1)**. Similarly, we assessed whether final colony size Gini coefficients were influenced by colony numbers per plate **(Supplementary Figure S4B)**.

In contrast to appearance time, colonies derived from 3- and 7-hour growth phases showed positive significant profiles, with weaker associations at 24- and 48-hour colonies, suggesting that colony size Gini coefficients at earlier growth phases should be interpreted with these differences in mind. Additionally, to understand the relationship between colony intensity and size, we compared the summed pixel intensity to the final colony size **(Fig. 2C)**. The results showed that colonies from 48 hours of growth displayed the highest variation in intensity per size. In contrast, this heterogeneity was less pronounced in the other growth phases. To assess the reproducibility of the *in vitro* distributions, the analyses presented in **Fig. 2A–C** were repeated for each biological replicate separately **(Supplementary Figure S5)**. The heterogeneity profiles observed in **Fig. 2A-C** across growth phases were preserved between replicates, with 48-hour sample is consistently showing the highest Gini coefficients across appearance time, final colony size, and intensity per size, despite minor variation across the biological replicates. Pairwise Spearman correlation analysis of appearance time distributions between replicates confirmed significant agreement across all growth phases (Spearman *ρ* ≥ 0.994 for all comparisons; **Supplementary Table S2**), indicating that the phenotypic heterogeneity is reproducible across independent experiments. While heterogeneity trends across growth phases followed a similar pattern for all three features, the absolute Gini coefficients differed systematically, appearance time and intensity per size representing lower Gini coefficients than final colony size **(Fig. 2A-C)**. This likely arises from the Gini coefficient’s dependence on the distribution mean. As appearance time and intensity per size values are positioned further from zero than colony size, an equivalent degree of spread yields a proportionally smaller Gini coefficient.

To further assess differences in appearance times, the complementary cumulative distribution function (1-CDF) of appearance times was calculated **(Fig. 2D)**, a previously established approach for characterizing heterogeneity in microbial population appearance times^11^. The results were reflected in distribution plots, where 3- and 48-hour samples showed broader and less steep profiles, indicating higher heterogeneity, while 7- and 24-hour growth periods displayed steeper curves with less heterogeneous appearance times. These patterns were summarised by the mean slope values **(Fig. 2E)**, which capture the steepness degree of heterogeneity across diverse growth periods. Quantifying heterogeneity through distributional shape (including tails and CDFs) is standard in microbial phenotyping and growth-variability studies^11^. To statistically quantify distributional differences in phenotypic features across growth phases, we calculated the Kolmogorov-Smirnov (KS) test and 90th percentile quantile differences for all pairwise condition comparisons **(Fig. 2F)**. The KS D-statistic measures the maximum divergence between two cumulative distributions, enabling comparison of appearance times or other morphologies across growth phases. Across all three metrics, 3 vs. 7 hours consistently produced the smallest D-values, while comparisons pairing an early timepoint with 48 hours of growth produced the largest **(Fig. 2F, upper panel)**. The 90th percentile quantile differences further highlighted appearance time differences at the right tail of the distribution, showing how far apart the slowest-growing colonies were between conditions, with 3 or 7 hour growth in comparison to 48 hour growth showing the largest positive shifts **(Fig. 2F, lower panel)**. Comparisons between 3 and 7 hours growth produced smaller values for both statistics, reflecting their close similarity. The same trend was observed for final colony size and intensity per size, with comparisons involving 48 hour showing the largest differences. For final colony size, this pattern was less consistent, with 24 vs. 48 hour growth showing a smaller shift. These results may suggest that colonies from 3 and 7 hours remain in comparable growth states, whereas colonies from 48 hour show distinct phenotypic distributions, consistent with the heterogeneity profiles observed in **Fig. 2A-C**.

### Appearance time relates differentially to colony morphology

To further assess whether colonies with early or late appearance times exhibited distinct morphological characteristics, we compared appearance time to colony size, intensity per size, circularity, eccentricity, and solidity. Colonies with earlier appearance times reached to larger size, whereas later appearing colonies showed smaller size, demonstrating a significant negative correlation between final colony size and appearance time across all growth phases (Spearman *ρ* ranging from -0.20 to -0.74) **(Fig. 3A)**. Early appearing colonies from 3 hours growth showed greater heterogeneity compared to the colonies of other growth phases, yet the negative association between final size and appearance time remained consistent. A similar trend was observed for intensity per size, which generally decreased as appearance time increased, particularly for the 3-hour (Spearman *ρ* = -0.73) and 48-hour (Spearman *ρ* = -0.83) cultures **(Fig. 3B).** Morphologies based on colony shape varied slightly between the early- or late-appearing colonies. The 7 hours growth showed a small increase in circularity with longer appearance times and rounder colonies, whereas the 3 hours growth showed a slight increase in eccentricity with longer appearance times and more elongated colonies **(Fig. 3C and 3D)**. *Y. pseudotuberculosis* colonies generally appear with a circular-like shape. Therefore, even a small increase in elongation, which can arise from the image quality and the colony details captured by our system, can lead to large changes in eccentricity, while circularity can be affected less and stay close to circle. Even though the variances were low, solidity values were reflected by the circularity trends across different growth phases **(Fig. 3E)**. Colonies generally exhibited high solidities, indicating overall smooth colony boundaries. Association between appearance time and downstream colony outcomes is consistent with studies showing that lag-time tuning shapes survival and growth under fluctuating stresses^13^. Finally, while a negative correlation between appearance time and final size is mathematically expected in a fixed-time assay, we sought to determine whether this was purely an artifact of delayed initiation or driven by inherently altered growth kinetics. By fitting the time-dependent increase in colony area to a Gompertz model^27^, maximum growth rates were computed independently of appearance time, we observed a wide distribution of growth rates *in vitro* **(Fig. 3F)**. While many colonies maintained similar maximum growth rates regardless of appearance time, others exhibited distinct kinetic trajectories. Notably, there was a negative correlation between appearance time and maximum growth rate, a phenotype that was particularly pronounced in the 48-hour samples (Spearman *ρ* = -0.555). This might indicate that later-appearing colonies do not merely have less time to grow but could exhibit slower maximum growth rates specifically at 48-hour samples (**Fig. 3F**). To test whether these results depend on the Gompertz model, each colony’s maximum growth rate was re-estimated without assuming a growth curve, as the maximum slope of the area trajectory *A*(*t*) over a sliding window of five consecutive timepoints. This model-free estimate agreed closely with the Gompertz model for individual colonies (Spearman *ρ* = 0.95–0.97 in all groups; **Supplementary Figure S6**). The negative association between appearance time and growth rate was preserved (e.g. 48-hour: *ρ* = −0.568 model-free vs. −0.555 Gompertz). This variance is consistent with colonies differing in post-initiation growth rate as well as in time to detection. At the present imaging resolution, however, the duration of a physiological lag phase and the early growth rate cannot be estimated independently, and appearance time should therefore be interpreted as an operational measure rather than a direct measure of lag.

**Fig. 3.**
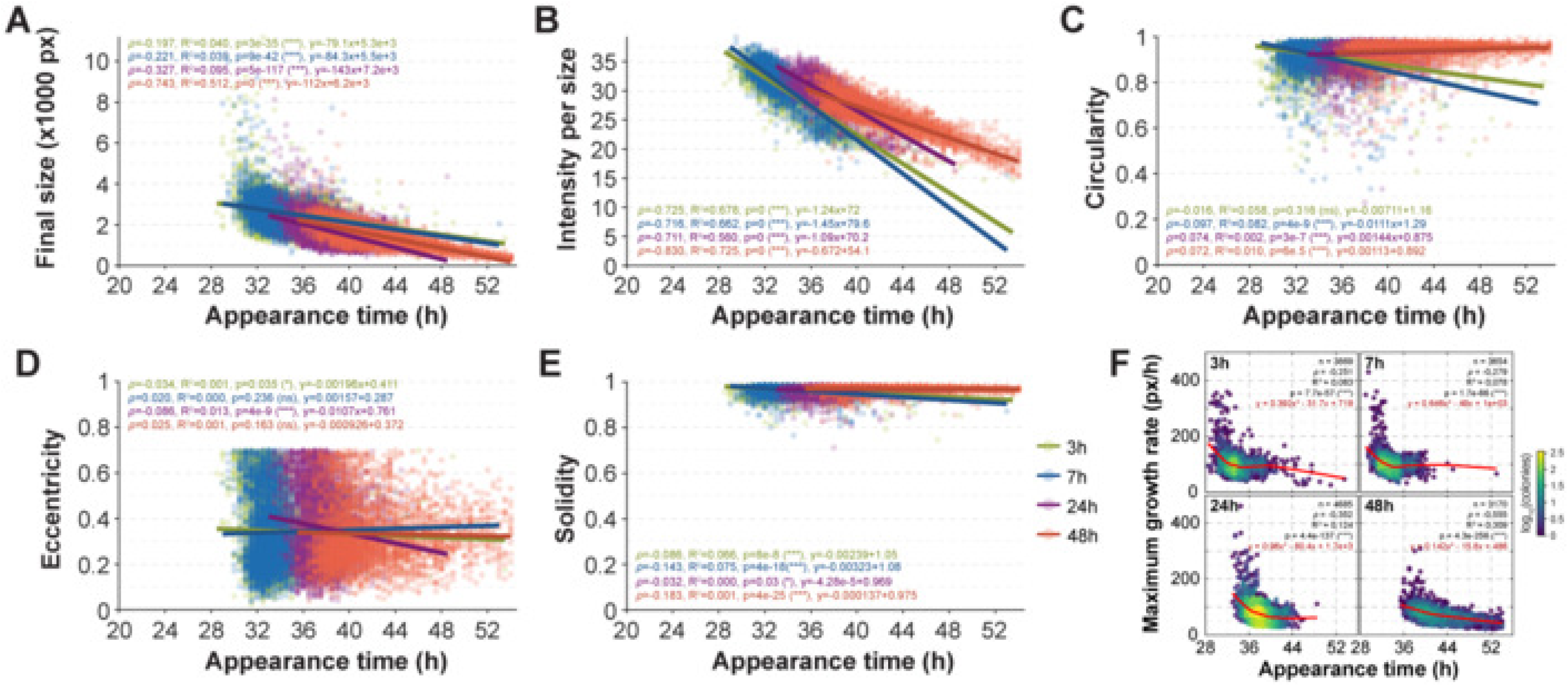
Comparison of colony morphology and appearance time related heterogeneity across growth phases. **A-E** Comparison of appearance times (x-axis, h) to colony morphologies (y-axis) for *in vitro* samples; final colony size **(A)**, intensity per size **(B)**, circularity **(C)**, eccentricity **(D)**, and solidity **(E)**. Points and matching darker lines are colored by growth time. Each point represents an individual colony. Lines represent the fitted trend for each sample group, where the slope of the line indicates the direction and strength of association between the appearance time and colony morphology. **F** Relationship between appearance time and the maximum growth rate per hour for individual colonies across growth conditions are represented in scatter plots. Growth rates were derived by fitting a Gompertz model to time-course area measurements. Points are colored by local colony density (log-scaled) to indicate the number of overlapping colonies, and red lines represent LOESS smooth fit. *n* represents the total number of colonies across all biological and technical replicates. **A-F** Spearman correlation coefficients (*ρ*), coefficients of determination (R^2^) corresponding *p*-values (*p*), and fitted linear equations (y = mx + b) are indicated for each growth condition in matching sample colors.

### HeteroTyper reliably quantifies phenotypic heterogeneity across diverse dark media

To determine whether HeteroTyper can reliably evaluate colony phenotypes on dark media prior to investigating heterogeneity in *in vivo*-derived bacteria, we tested the system on three types of dark agar media on 48-hour grown cultures, exhibiting the most heterogeneity. *Y. pseudotuberculosis* cultures were inoculated onto Blood Agar (BA), MacConkey (MC), and YSA, and colony phenotypes were assessed using HeteroTyper. Although BA and MC are not optimal growth media for *Yersinia*, they were deliberately selected to induce varying metabolic states and distinct population structures. Automated colony counts showed strong concordance with manual counts across all three media types, demonstrating the robustness of segmentation across tested dark agar media **(Supplementary Figure S7A)**. Crucially, HeteroTyper could capture the divergent growth kinetics and varying degrees of heterogeneity driven by each environment. Bacteria grown on BA and YSA displayed rapid growth and broad phenotypic distributions in appearance time and final colony size (Gini = 0.097 and 0.071, respectively). Notably, YSA cultures also exhibited the highest heterogeneity in colony intensity per size, highlighting the system’s sensitivity to subtle optical and morphological variations. Conversely, the selective pressures and suboptimal conditions of MC medium imposed metabolic constraints that resulted in prolonged lag phases, restricted colony sizes, and a markedly more uniform population overall (Gini = 0.044) (**Supplementary Figure S7B–E**). Overall, these results demonstrate that HeteroTyper can effectively track colonies and quantify phenotypic features across diverse dark media, establishing a robust baseline for subsequent analysis of *in vivo*-derived bacterial populations.

### Heterogeneity in appearance time reveals diverse growth phenotypes in infected tissues

To assess whether phenotypic heterogeneity was present in bacteria recovered from mouse infection, we analysed the colony phenotypes of *Y. pseudotuberculosis* cells isolated from different organs including MLN, spleen, and liver. As number of colony-forming units cannot be accurately estimated and are highly variable among organs and individual mouse prior to plating, we screened 10^2^-10^5^ dilution series on dark YSA plates and selected the plates with appropriate colony numbers (10-500 colonies). However, for some of the organs, specifically liver, the number of colonized bacterial cell were very low leading lower bacterial colonies screened with HeteroTyper. Nevertheless, considering the Gini coefficients, the appearance time distribution was observed to be more heterogeneous for colonies recovered from MLN (Gini = 0.059) compared to spleen (Gini = 0.04) and liver (Gini = 0.043), reflected in the wider spread observed in the appearance time probability density plots **(Fig. 4A)**. On the other hand, appearance time appeared generally more heterogeneous in colonies recovered from infection in comparison to *in vitro* grown bacteria **(Fig. 4A and 2A)**. Interestingly, colonies originating from the MLN were generally smaller, with higher heterogeneity (Gini = 0.207) compared to that of the spleen (Gini = 0.133) and the liver (Gini = 0.160) **(Fig. 4B)**. We repeated the same pairwise hypothesis testing framework used for **Fig. 2**, confirming that colony size differed significantly between the MLN and both the spleen and liver (Wilcoxon rank-sum, *p* < 0.02 for both; **Supplementary Table S3**).

**Fig. 4.**
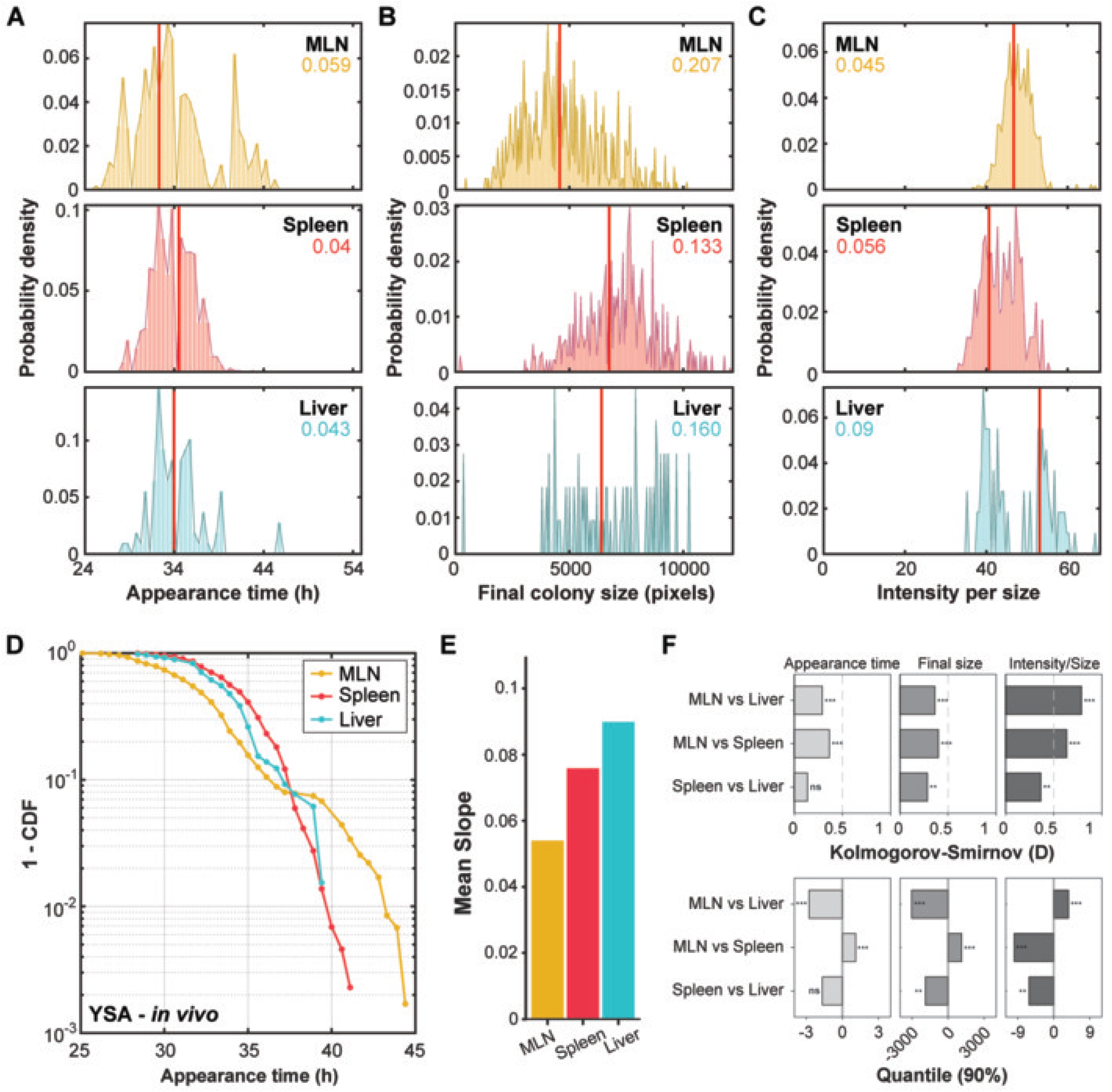
Host microenvironments drive divergent appearance time profiles across organs. For *in vivo* experiments, FVB\n mice were orally infected with wt *Y. pseudotuberculosis* YPIII. At 5 days post-infection, MLN, spleen, and liver were collected, homogenized, serially diluted, and 10^2-5^ dilutions were plated on YSA plates, incubated at room temperature for 24 hours and imaged in HeteroTyper system. **A** Appearance time, **B** final colony size, and **C** summed pixel intensity per colony size distributions using probability distribution function of bacterial colonies on agar plates originating from cells grown for MLN (n = 663), spleen (n = 469), and liver (n = 88). **D** Appearance times of *in vivo* (plated on YSA) cells are plotted as complementary cumulative distribution function (1–CDF) on logarithmic scale. **E** Mean slope values of each sample group represented in **D**. **F** Pairwise comparisons between organ types are represented for appearance time, final colony size, and intensity per size using bar plots. Kolmogorov–Smirnov statistics (D) are displayed in the upper panel, and differences in the 90th percentile are shown in the lower panel. Appearance time is represented in hours (h), with an additional incubation period of 24 hours, while the colony size was measured in pixels (px). For each parameter in **A-C**, Gini coefficients are shown in the colors of organ type and median values are indicated by red vertical lines. Data presented represent pooled colonies recovered from all animals that yielded sufficient number of colonies for quantification: MLN (n=3 mice), Spleen (n=3 mice), and Liver (n=2 mice). Asterisks indicate statistical significance (*p* < 0.05: *, *p* < 0.01: **, *p* < 0.001: ***, ns: not significant) in **F**.

Notably, the number of colonies recovered from liver were relatively lower than bacteria recovered from other organs, likely because the liver is colonized later during the systemic infection of *Y. pseudotuberculosis*, resulting in lower bacterial burden. Nevertheless, these numbers of colonies may not be sufficiently enough to reach such conclusion of least heterogeneity in liver. It’s also noteworthy that the tail of the appearance time distribution for liver indicate that there could be similar degree of heterogeneity in liver too **(Fig 4A)**. In contrast, colonies from the MLN exhibited higher intensity per colony size than those from the spleen, whereas colonies from the liver generally showed the highest intensity per colony size among all organs, although the number of liver-derived colonies was very low **(Fig. 4C)**. The intensity per size feature displayed a Gini coefficient of 0.045 for the MLN. Cumulative appearance time distributions were also calculated for *in vivo* samples **(Fig. 4D)**. Colonies from MLN exhibited the least steep profile, indicating higher heterogeneity, while the spleen showed the steepest curve, consistent with the degree of heterogeneity and Gini coefficients observed in the distribution plots **(Fig. 4D and 4A)**. These results were consistent with mean slope values **(Fig. 4E)**.

To rigorously validate these differences despite varying sample sizes, we performed pairwise KS tests and computed 90% quantiles for each feature **(Fig. 4F).** The **KS D-statistics** confirmed that the MLN appearance time distribution is statistically distinct from both the spleen and liver, characterized by a significantly wider spread. Furthermore, the **90% quantile** analysis highlighted that the MLN contains a prominent subpopulation of long-lagging cells, which is significantly more pronounced than in the peripheral organs **(Fig. 4F)**. These statistical measures, combined with the Gini coefficients **(Fig. 4A–C)**, suggest that the MLN promotes a unique level of phenotypic heterogeneity **(Fig. 4F).** Organ- and niche-specific phenotypic diversification during infection is well documented in enteric pathogens and supports the interpretation of stable, tissue-structured subpopulations^2,3^.

### Host microenvironments shape appearance time and colony morphology during infection

To investigate how host microenvironments influence morphologies of colonies with varying degree of appearance time, we analysed colonies recovered from different organs. Generally, the final colony size of colonies derived from infection was negatively associated with appearance time, similar to the *in vitro* results **(Fig. 5A)**. MLN displayed greater size heterogeneity than spleen and liver, particularly among early-appearing colonies **(Fig. 5A)**. Intensity per size also declined as appearance time increased, and colonies with similar appearance times were more diverse in liver than in MLN and spleen **(Fig. 5B)**. Circularity phenotypes differed by organ, with MLN colonies appearing more circular, liver colonies resembling MLN, and spleen colonies generally less circular **(Fig. 5C)**. Eccentricity was generally similar across organs with only minor variations **(Fig. 5D)**. Solidity remained high overall and reflected the circularity phenotypes, as also observed in *in vitro*. Colonies from MLN showed higher solidity reflecting smoother colony edges, whereas spleen showed lower solidity indicating more irregular colony boundaries **(Fig. 5E)**. These organ-resolved phenotypes align with reports that within-host heterogeneity reflects differential stress histories and microenvironments^2,3^. Moreover, colonies recovered from the MLN showed no significant association between maximum growth rate and appearance time (Spearman *ρ* = 0.012, *p* = 0.76), while colonies derived from spleen showed a weak but significant relationship (Spearman *ρ* = -0.138, *p* = 2.77e-3), suggesting that appearance time variability in these tissues is not primarily driven by maximum growth rate **(Fig. 5F)**. Colonies derived from the liver exhibited a moderate positive association (Spearman *ρ* = 0.380, *p* = 2.65e-4), though the relatively low colony numbers across all infected tissues suggest that these relationships should be interpreted with caution.

**Fig. 5.**
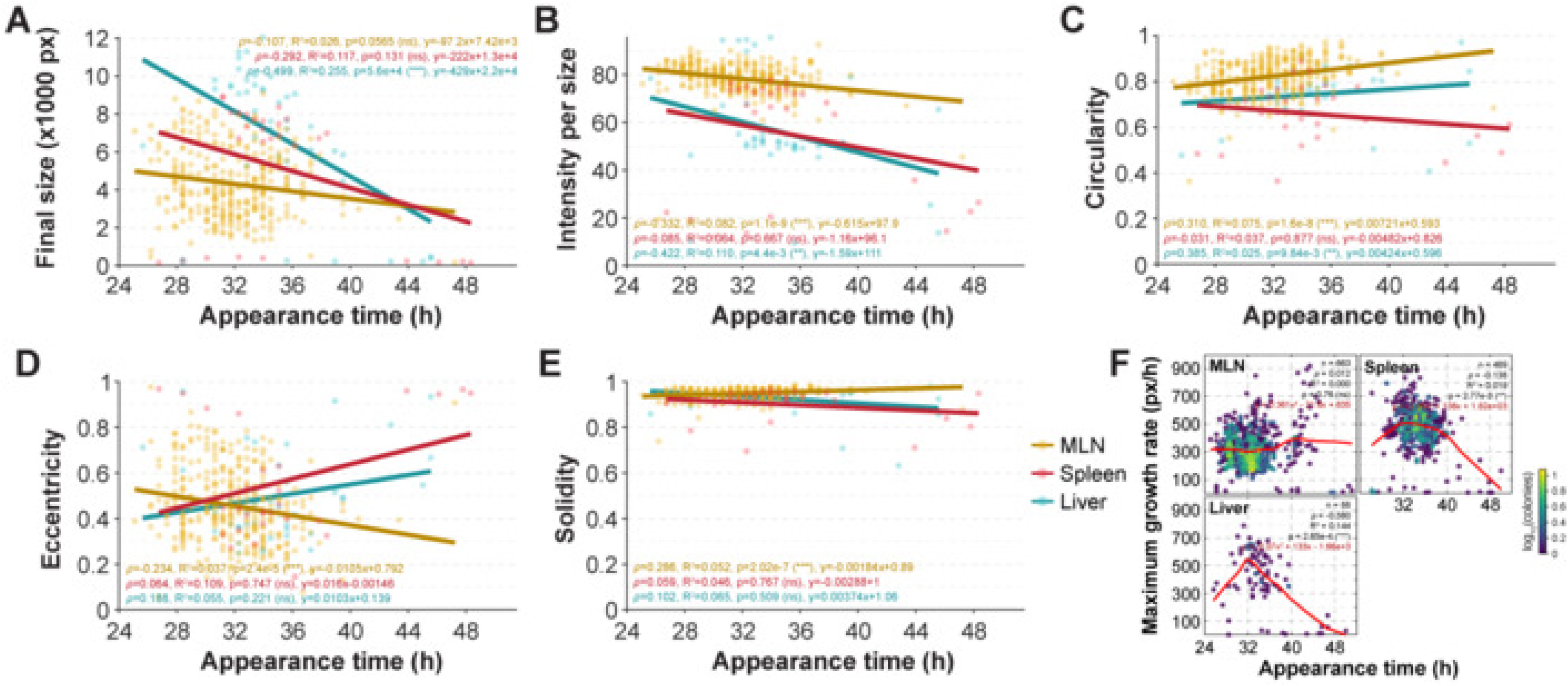
Comparison of colony morphology and appearance time related heterogeneity across host compartments during infection. **A-E** Comparison of appearance times (x-axis, h) to colony morphologies (y-axis) for *in vivo* samples; final colony size **A**, intensity per size **B**, circularity **C**, eccentricity **D**, and solidity **E**. Points and matching darker lines are colored by organ type. Each point represents an individual colony. Lines represent the fitted trend for each sample group, where the slope of the line indicates the direction and strength of association between the appearance time and colony morphology. **F** Relationship between appearance time and the maximum growth rate per hour for individual colonies across growth conditions are represented in scatter plots. Growth rates were derived by fitting a Gompertz model to time-course area measurements. Points are colored by local colony density (log-scaled) to indicate the number of overlapping colonies, and red lines represent a LOESS smooth fit. *n* represents the total number of colonies across all biological and technical replicates. **A-F** Spearman correlation coefficients (*ρ*), coefficients of determination (R^2^) corresponding *p*-values (*p*), and fitted linear equations (y = mx + b) are indicated for each growth condition in matching sample colors. Data presented represent pooled colonies recovered from all animals that yielded sufficient number of colonies for quantification: MLN (n=3 mice), Spleen (n=3 mice), and Liver (n=2 mice).

## Discussion

The variability in appearance time among bacterial cells in a genetically uniform population has been associated with prior exposure to stress conditions and with extended periods of growth^22,28,29^. Such variability may reflect both the magnitude of the stress experienced and the heterogeneity of stress responses among individual cells. In line with these observations, we observed the largest cell-to-cell variation in colony appearance times for *Y. pseudotuberculosis* populations derived from extended growth. In addition, we detected a negative correlation between appearance time and colony intensity normalized by size, which may indicate that colonies appearing late tend to have lower density. Therefore, we speculate that this pattern could reflect a link between prolonged ppearance time, slower growth, and altered quorum sensing or motility behaviors^30,31^, although this remains to be tested. If so, it would suggest that appearance time is not only a survival trait but may also be coupled to downstream growth dynamics at the colony level, which requires experimental validation.

Bacterial phenotypic heterogeneity is expected to be even more pronounced during infection, where colonization sites are characterized by complex and spatially structured microenvironments containing gradients of nutrients, oxygen, pH, antimicrobial factors, and immune effectors. Such heterogeneity provides a fitness advantage by enabling subsets of the population to better adapt to local conditions, evade or subvert immune-mediated killing, and tolerate antimicrobial treatment^32-37^. Our understanding of how such microenvironmental heterogeneity is organized within infected tissues, and how it shapes the degree of phenotypic heterogeneity required for long-term survival and treatment failure, is still limited. To address this, we quantified single-cell appearance time distributions of *Y. pseudotuberculosis* cells recovered from MLN, spleen, and liver of orally infected mice. We found the highest appearance time heterogeneity among *Y. pseudotuberculosis* cells isolated from MLN. Following oral infection, *Y. pseudotuberculosis* crosses the intestinal barrier, colonizes Peyer’s patches and the intestinal mucosa, and then disseminates to MLN, from which it can spread further to the spleen and liver^38-42^. MLN are populated by diverse immune cell subsets and, during *Yersinia* infection, show robust infiltration of neutrophils, natural killer (NK) cells, monocytes, and macrophages, as well as sustained tissue remodelling and chronic lymphadenopathy^38,43^. Such an immune-rich and dynamically changing niche likely generates strong spatial and temporal heterogeneity in stresses encountered by individual bacteria, promoting the emergence of subpopulations in distinct physiological states with divergent appearance times. By contrast, *Y. pseudotuberculosis* cells recovered from spleen and liver displayed narrower appearance time distributions, indicating reduced phenotypic variation at these sites. One possible explanation is that by the time bacteria successfully colonize these secondary organs, strong population bottlenecks and selective pressures in upstream tissues (intestine and MLN) may already have filtered out many phenotypic variants, resulting in a more homogeneous population with similar physiological states. Alternatively, the spleen and liver microenvironments may impose more uniform stress conditions, leading to tighter control of growth resumption dynamics. Together, our findings support a model in which *Y. pseudotuberculosis* adapts to infection-site– specific microenvironments by modulating the breadth of phenotypic heterogeneity in appearance time, with particularly pronounced phenotypic heterogeneity in the MLN. Colonies derived from infected tissue showed much greater variation in appearance time and maximum growth rate than colonies derived from *in vitro* cultures. One likely explanation is that bacteria in tissue are spread across different microenvironments and are therefore exposed to different spatial conditions, such as nutrient availability, oxygen levels and host immune pressure. In contrast, *in vitro* cultures are exposed to a relatively uniform environment. These environmental cues may therefore shape both how long bacteria take to resume growth and how fast they grow afterwards. Notably, colonies from the MLN included a subgroup that appeared late but reached a higher maximum growth rate. This combination of late appearance and fast subsequent growth is of particular interest in the context of antibiotic persistence, where lag time and growth rate have so far been considered largely independent traits. Further work is needed to determine whether this subpopulation is more tolerant to antibiotics.

Overall, these results illustrate that bacterial adaptation during infection is both organ-specific and dynamically shaped by tissue microenvironments and host immunity. The technical improvements in HeteroTyper combined with the computational pipeline provides a scalable framework to quantify single-cell appearance time distributions in *in vitro* settings and infection. This approach will facilitate future studies dissecting the mechanisms by which phenotypic heterogeneity of bacterial pathogens and may help to develop novel treatment strategies and fight against antibiotic resistance.

While HeteroTyper demonstrates robust performance for *Y. pseudotuberculosis*, several technical limitations require attention. Current limitations include operation at room temperature, which constrains applicability to pathogens with alternative temperature requirements; dependence on circular colony morphology; and approximately 20% ticker medium in a plate to prevent agar cracking. Despite these constraints, ongoing developments including temperature-controlled incubation and adaptive morphological recognition will extend the platform’s applicability to a broader range of clinically relevant bacterial species.

## Materials and Methods

### Image acquisition of bacterial populations

HeteroTyper system uses a GoPro Hero 10 Black camera to track the growth of bacterial colonies on agar plates. The center coordinates for the camera that was attached to the cutting head of the modified laser cutter, HeteroTyper, were set with the LightBurn software (LightBurn Software v. 1.3.01, LLC, 2022). These coordinates were aligned with center areas of each of the 104 plate holders to acquire images of each agar plate placed to plate holders in HeteroTyper. The cutting head with the camera attached was mounted on the guide rail of the laser cutter, allowing it to move along both the x and y axes to capture images of the plates from above. The time lapse images were taken in 30-minute intervals, which were set via the potentiometer on the Arduino Nano board connecting laser cutter and camera via its remote control. The signal from laser cutter was transmitted from the Arduino Nano board to the remote control at the end of each time interval to trigger the camera to capture an image of each plate. The agar plates were imaged while the lid was open. The images were collected in .jpg file format on a SD card placed in the camera, transferred to a computer, and grouped into separate folders for each sample group using the metadata information with R programming^44^.

### Computational pipeline for colony screening

The computational pipeline used in analysis was developed in MATLAB 2023b (The Math Works, I.)^45^. Preliminary efficiency tests were performed on a computer with Windows 10, i7 core, and 16 GB RAM. The analysis of the extended growth and stress exposure experiments in this study were performed on a computer with Windows 11, i9-13980HX 2.20 GHz, and 64 GB RAM using MATLAB 2023b version.

### Colony detection and segmentation

The computational pipeline began with loading the raw images and metadata information to MATLAB. The parameters of the images were defined, such as border range, minimum colony size in pixels (px), minimum and maximum colony numbers on the plates, intensity threshold, and eccentricity threshold. Following this, plate centers were automatically detected for each plate individually. A subset of frames (five, by default) from each plate’s own image stack were sampled, with center and radius estimated per frame via Hough-transform circle detection, followed by calculation of a median center value. Plates with centers deviating by more than 10% from this median were highlighted with a red circle to indicate potential alignment errors, while correctly aligned plates were marked in green. A summary figure with all plates was displayed, enabling visual verification of correctly detected plate centers. Accurate plate alignment is critical for consistent downstream quantification. Therefore, plates that did not fit the requirement for the center deviation, minimum, and maximum colony numbers were excluded from the analysis.

Colony detection and segmentation were performed using the last image in each time-lapse stack. The images were initially converted to grayscale to simplify image data, and uneven lighting across the plate was reduced by using a Gaussian filter (see Online Methods and https://github.com/avicanlab/HeteroTyper). Gaussian-corrected images were subtracted from the original image, which minimized the light differences between the center and the edges of the plate. The intensity values of the background-corrected image were normalized to a 0-1 scale based on the minimum and maximum pixel intensities in each image, which can reduce variations caused by lighting differences between plates. The contrast of the images was slightly adjusted, and a small Gaussian blue was applied to smooth minor intensity variations and reduce background noise without altering colony shapes.

Colonies were then identified by applying intensity threshold to the smoothed image, which separates darker colonies from the brighter background. Small background artifacts below the defined minimum size (100 px) were removed, as objects of this size or smaller likely noise or dust particles rather than true colonies. This threshold was determined by inspecting several representative images and selecting a value that can minimize false detections. Small gaps within detected colonies were filled to prevent fragmentation of single colonies into multiple smaller colonies. Colonies that were touching or partially overlapping were separated using a marker-controlled watershed, with markers placed at each colony’s own interior peak rather than a fixed erosion radius or centroid, splitting colonies of different sizes appropriately.

Colonies located near the plate edges were removed using a circular mask corresponding to approximately 85% of the plate radius, in which helped to avoid reflections, possible agar cracks or partial colony detections near the border but may have excluded a few valid colonies. A size threshold of 100 px was applied to exclude false detections, as these are likely background noise rather than positive colonies, together with an eccentricity filter of 0.70, which colonies above this threshold are removed to exclude non-circular or merged colonies. Colonies whose centroid fell within a narrower zone near the border were subjected to a stricter eccentricity threshold of 0.55, since partially cropped or overlapping colonies near the plates’ detection edge are likely to appear elongated.

To evaluate the scalability of the HeteroTyper across different experimental conditions, colony detection was performed on both transparent Luria-Bertani Agar (LA) plates used for *in vitro* and dark YSA plates were used for *in vivo* samples. The segmentation approach was applied globally, using the same parameters across different media types, without requiring media-specific adjustments. Shape or intensity-related phenotypic parameters, such as colony size, total pixel intensity, perimeter, solidity, circularity, and eccentricity, were measured for all colonies that has passed the segmentation. These measurements were used to generate growth profiles by tracking colony size and summed pixel intensity over time. Final colony size and appearance time were obtained from these measurements, where appearance time corresponded to the first time point at which a colony reached at least 100 px in size.

### Gini coefficient

Phenotypic heterogeneity within a population was quantified using the Gini coefficient (G), which measures how unequally a feature is distributed among colonies. For n colonies with values X^1^ to X^n^, G is the mean absolute difference between all pairs of colonies divided by twice the mean value:

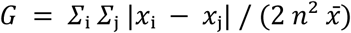

G ranges from 0 to 1. A Gini coefficient of 0 indicates that all colonies have the same feature value, and higher values indicate greater diversity among colonies relative to the group mean. For variables that do not approach zero, such as appearance time measured from inoculation, the value of G is lower than it would be if a zero-based time origin were used. This occurs because adding a constant increases the mean without altering the differences between colonies. Therefore, G was compared only between groups analysed using the same time origin and measurement units.

G was computed on unbinned colony values using the sorted-value form *G* = 2 ∑ *i* · *x*(i) / (*n* ∑ *x*) − (*n* + 1)/*n*, where *x_(i)_* is the *i*-th value in ascending order. Colonies were included if their final area was at least 100 px and their appearance time was not censored. The estimator was not corrected for sample size, so its maximum is *(n − 1)/n*.

### Maximum growth rate (Gompertz) analysis

To estimate the maximum growth rate (*μ*max), the colony area over time was fitted for each colony. Colony area values were smoothed with a three-point median filter, and only colonies with at least six positive, finite area measurements were fitted. Each colony’s growth curve was fitted to a Gompertz model:

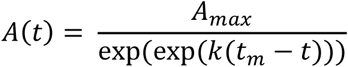

where *A*(*t*) is colony area (px) at time *t*, *A*max is the maximum colony area, *k* is a rate constant (h^-1^) describing how steeply colony area increases, and *t*m is the time (h) at which growth is fastest. The maximum growth rate of each colony was computed from the fitted parameters as

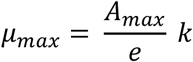

where *e* is Euler’s number (≈ 2.718), and is reported in pixels per hour (px/h).

Fitting was performed for each colony in MATLAB by nonlinear least squares (lsqcurvefit, maximum of 500 iterations, function tolerance 1e-8), using only time points with positive, finite area values. Starting values were the largest observed area for *A*_max_, 0.3 h^-1^ for *k*, and the time of the steepest observed increase in area for *t*_m_. These starting values served only as initial guesses, and the final values were estimated separately for each colony. Parameters were restricted to realistic ranges: *A*max could not be negative, *t*m must lie within the imaging period, and *k* was limited to 0-5 h^-1^ to exclude biologically impossible, near-instantaneous increases in area. A fit was accepted when the optimizer converged within these bounds. No additional fit thresholds were applied. Colonies for which the fitting failed, which had fewer than six measurements, or which had no defined appearance time were excluded from growth-rate analyses.

For the remaining colonies, the relationship between *μ*max and appearance time was assessed with a two-sided Spearman rank correlation (*ρ*, *p*-value) per sample group. The R^2^ are shown as *ρ*^2^. Each colony is coloured by the log^10^ number of colonies in its local density bin (a 40×40 bin grid over appearance time and *μ*_max_). The overall trend was visualised with a locally estimated scatterplot smoothing (LOESS) curve, using local linear regression with tricube distance weights and a span of 0.75, so that nearest 75% of colonies along the appearance time axis contribute to each local fit. No robustness iterations were performed, and the curve was constrained to non-negative values within the displayed growth-rate range. The equation shown on each figure panel is a second-degree polynomial fitted to the LOESS curve to summarise its shape. The LOESS curve, local density and polynomial equation was used for visualisation only and were not used for inference.

### Statistical analysis

Phenotypic heterogeneity within each plate or group was quantified using the Gini coefficient, calculated on raw per-colony values for appearance time and final colony size, and correlated against colony count per plate (Pearson and Spearman) to assess potential bias from colony density. Distributional differences were further characterized using the Kolmogorov-Smirnov (KS) D statistic and the difference in 90th-percentile (Q90) values between groups, capturing shifts in the upper tail. Pairwise concordance between biological replicates within each experimental group was assessed on quantile-binned growth profiles using Pearson and Spearman correlation, applied identically across all *in vitro* and *in vivo* experimental groups. The relationship between appearance time and individual morphological features (colony size, eccentricity, solidity) was assessed per condition using Pearson correlation (R²) and Spearman rank correlation (*ρ*), with linear fit equations annotated on scatter plots, and significance markers were based on Spearman *p*-values for robustness to non-normality and outliers. Pairwise comparisons between experimental conditions were performed for each feature using a normality-aware two-sample test. Groups were first assessed for normality via the Lilliefors test, followed by Wilcoxon rank-sum test as distributions were not normally distributed, with resulting *p*-values corrected using the Benjamini–Hochberg false discovery rate (FDR). For the maximum growth rates the relationship between *μ*max and appearance time was assessed using Spearman rank correlation (*ρ*), with R² reported as *ρ*².

### Visualization of bacterial colonies on agar plates

Various function packages in MATLAB were employed for the visualization of the data. Raw and analysed plate images for plate center identification and segmentation were printed on screen using “figure” function. This function was also used to combine the plots for distinct growth features in a single window. The “plot” function was used to generate line graphs of growth curves. Comparisons between colony features were visualized as scatter plots, including density-colored scatter plots by using a custom “dscatter” function rather than the “plot” function. The “histogram” function was used to generate frequency bar plots for final colony size and intensity per size, and appearance time distributions. Adobe Illustrator v30.8.2 was used to improve the graphical context of the figures (Adobe Inc.).

### *Y. pseudotuberculosis* strains and growth conditions

The kanamycin-resistant wt *Yersinia pseudotuberculosis* (YPIII/pIBX) strain was used for both *in vitro* and *in vivo* experiments. Unless stated otherwise, the wt strain was grown in Luria-Bertani (LB) medium (BD Difco^TM^ LB Agar, Lennox) supplemented with 50 µg/mL kanamycin (LB+Km50).

For *in vitro* experiments, the wt strain was subcultured in LA supplemented with kanamycin (50 µg/mL). The wt strain was cultured overnight at 26°C and 160 rpm in LB. The overnight culture was co-cultured in the morning to a starting optical density at 600 nm (OD^600^) of 0.05 and grown in LB at 26°C and 160 rpm. Samples were collected at different time points (3, 7, 24, and 48 hours) during growth. OD^600^ values were measured at each time points. Collected cultures were then serially diluted in LB medium, and 100 µL of dilutions (from 10^-4^ to 10^-6^ cells) were spread onto LA plates, The plates were incubated at room temperature (RT) for 20 hours and imaged using the HeteroTyper system. Equal number of images per sample group was used during image processing to ensure consistency across time points.

To evaluate HeteroTyper performance across multiple dark media, the wt strain was cultured under identical conditions, with 48-hour *in vitro* cultures of *Y. pseudotuberculosis* inoculated onto three distinct media (Blood Agar, BA; MacConkey, MC; YSA). BA and MC plates were supplemented with 50 µg/ml Kanamycin, while YSA was supplemented with CIN (Cefsulodin, Irgasan, Novobiocin) antibiotic complex. Experiments were performed in three independent biological replicates. Plates were incubated for the same duration as the *in vitro* assays and subsequently imaged using the HeteroTyper system.

### Mouse infection

Mice were housed in accordance with the Swedish National Board for Laboratory Animals guidelines. All animal experiments were approved by the Animal Ethics Committee of Umeå University (Dnr A65-15). All experiments were performed using female mice, as this sex is the established standard for *Yersinia* infection models in our laboratory. While the potential influence of biological sex on colony-level phenotypic heterogeneity has not been extensively characterized, we utilized a single sex to minimize experimental variables and ensure consistency across within-host microenvironments. For infection, the bacteria were grown overnight in LB at 26°C, and concentrations were measured by absorbance at OD^600^. Cultures were resuspended to 10^7^ colony forming units (CFUs)/mL in sterilized tap water supplemented with 150 mM NaCl. The infection dose was then determined by viable count and drinking volume. Eight-week-old four FVB\n (Taconic Farms, Inc.) mice were deprived of food and water for 16 hours before oral infection with ∼10^7^ CFUs of wt *Y. pseudotuberculosis* YPIII strain, which were supplied in their drinking water for 6 hours. Mice were inspected frequently for signs of infection and to ensure that infected mice showing prominent clinical signs were euthanized promptly to prevent suffering. At 5 days post-infection (dpi), mesenteric lymph nodes (MLNs), spleens, and livers were dissected and homogenized in phosphate-buffered saline (PBS). Homogenates were serially diluted in PBS and plated onto YSA plates. The plates were incubated at RT for 24 hours and imaged using the HeteroTyper system to monitor colony growth.

## Code and data availability

The MATLAB code for the HeteroTyper pipeline is available as open-source software on GitHub (https://github.com/avicanlab/HeteroTyper), with comprehensive documentation hosted on ReadTheDocs (https://heterotyper.readthedocs.io/). The *in vitro* data generated in this study are deposited on Zenodo (https://doi.org/10.5281/zenodo.21163063).

## Acknowledgement

This work was supported by Swedish Research Council (No. 2021-02466), Kempestiftelserna (JCK22-0017), and the Medical Faculty at Umeå University (FS 2.1.6-281-22) to K. Avican, by Swedish Research Council Excellence Center grant (No. 2022-06543) for the Center for Modeling Adaptive Mechanisms in Living Systems Under Stress to K. Avican. We acknowledge µNordic Single Cell Hub (µNiSCH) and Small Animal Research Imaging Facility (SARIF) at Umeå University. K. Kochanowski is supported by the Spanish Ministry of Research and Innovation (Grant number: RYC2021-033035-I /AEI/10.13039/501100011033).

## Authors contributions

SGS, TD, NH, and KA conceived, designed, and performed the experiments and SGS, KK, and KA developed the pipeline and analysed the data. SGS and KA wrote the manuscript with input from TD, NH, and KK.

