## Supplementary Figures S1-S7 for "High-throughput, single-cell-encoded colony imaging reveals infection-linked heterogeneity"

[1]Department of Molecular Biology, Umeå University, Umeå, 90187, Sweden
[2]Integrated Science Lab (IceLab), Umeå University, Umeå, 90187, Sweden
[3]Umeå Centre for Microbial Research (UCMR), Umeå University, 90187, Umeå, Sweden
[4]Science for Life Laboratory, Umeå University, Umeå, 90187, Sweden
[5]Unitat mixta d'Investigació IRTA-UAB en Sanitat Animal. Centre de Recerca en Sanitat Animal (CReSA). Campus de la Universitat Autònoma de Barcelona (UAB), Bellaterra, 08193, Catalonia, Spain.
[6]IRTA. Programa de Sanitat Animal. Centre de Recerca en Sanitat Animal (CReSA). Campus de la Universitat Autònoma de Barcelona (UAB), Bellaterra, 08193, Catalonia, Spain.

**Supplementary Figure S1**

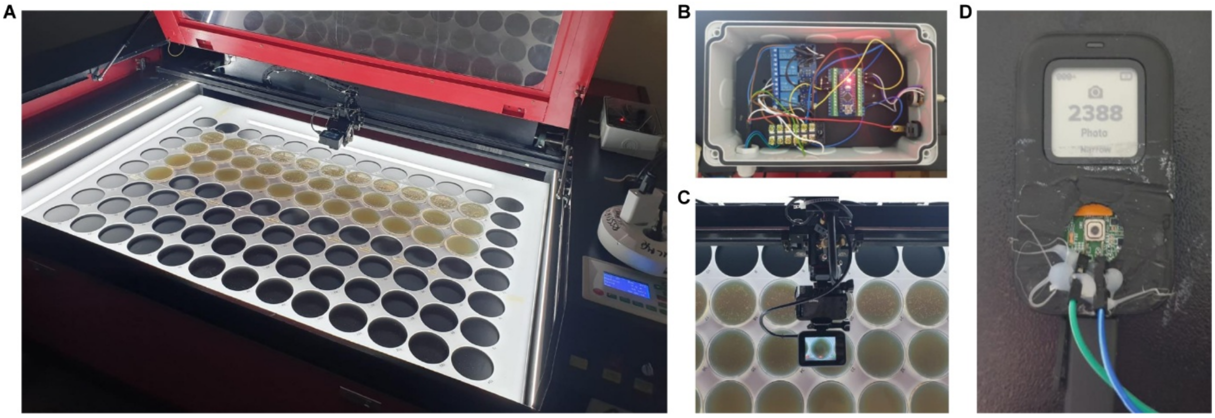


**Supplementary Figure S1. Setup of the HeteroTyper system.** The HeteroTyper system allows to grow bacteria on agar plates while simultaneously capturing images. **A** View of HeteroTyper from above while the lid is open. **B** Customized Arduino Nano board that receives a trigger signal from the laser-cutter and relays it to the remote control. **C** Go Pro Hero 10 camera mounted on the gantry and positioned above the plates for sequential imaging. **D** Remote control wired to the Arduino Nano to trigger the camera for automated imaging.

### Supplementary Figure S2

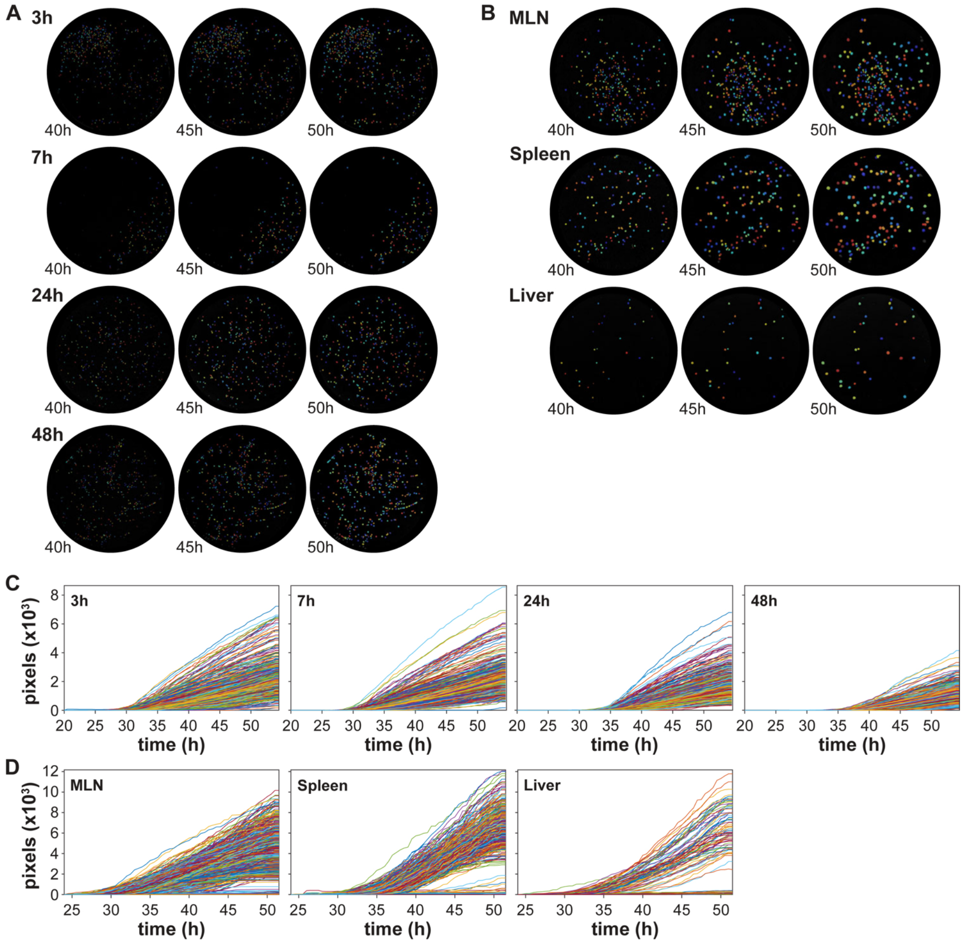


**Supplementary Figure S2. Time-lapse segmentation images and growth curves of *in vitro* and *in vivo Y. pseudotuberculosis* samples.** Segmentation images illustrating segmentation differences over time (40, 45, and 50 hours) for selected plates with colonies grown on LA plates **(A)** at different growth phases (3, 7, 24, and 48 hours), or on YSA plates **(B)** from bacterial cells recovered from mouse organs (MLN, spleen, and liver). Colors represent individual colonies identified. Growth curves of bacterial colonies on LA plates **(C)** correspond to the same growth phases shown in **A**, while growth curves of colonies on YSA plates **(D)** correspond to the same mouse organs shown in **B**. Colony size was measured in pixels and plotted against time (hours). Differences in the initial time points of growth curves reflect the starting point of imaging, which began before colonies became visible by naked eye and plates were incubated at room-temperature until the initial time point. **C-D** Data is combined for all the replicates for each sample group, with each line representing an individual colony.

### Supplementary Figure S3

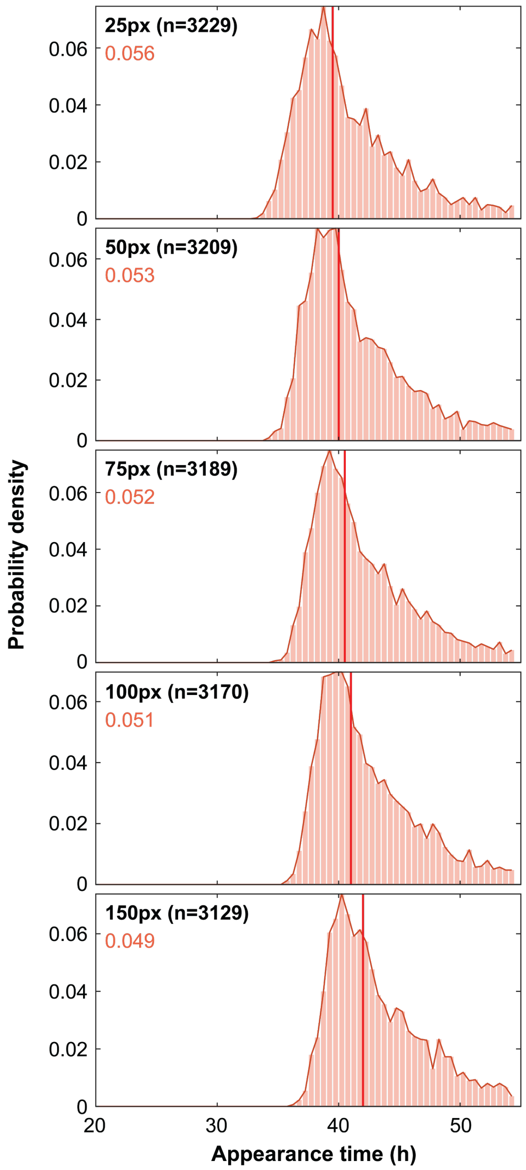


**Supplementary Figure S3. Colony appearance time distributions analysed using different colony size thresholds**. Probability density functions of appearance times are shown for *Y. pseudotuberculosis* colonies originating from cells harvested at 48 hours of growth. Colony appearance time was defined as the time a colony first reached a size threshold of 25, 50, 75, 100, or 150 pixels (px). The same plates and images as in Fig. 2 were re-processed at each threshold, except 100 px, and the panels are displayed from top to bottom in order of increasing threshold. Gini values are shown in the color assigned to 48 hours of growth, and median values are indicated by red vertical lines. n represents the number of colonies identified at each threshold.

### Supplementary Figure S4

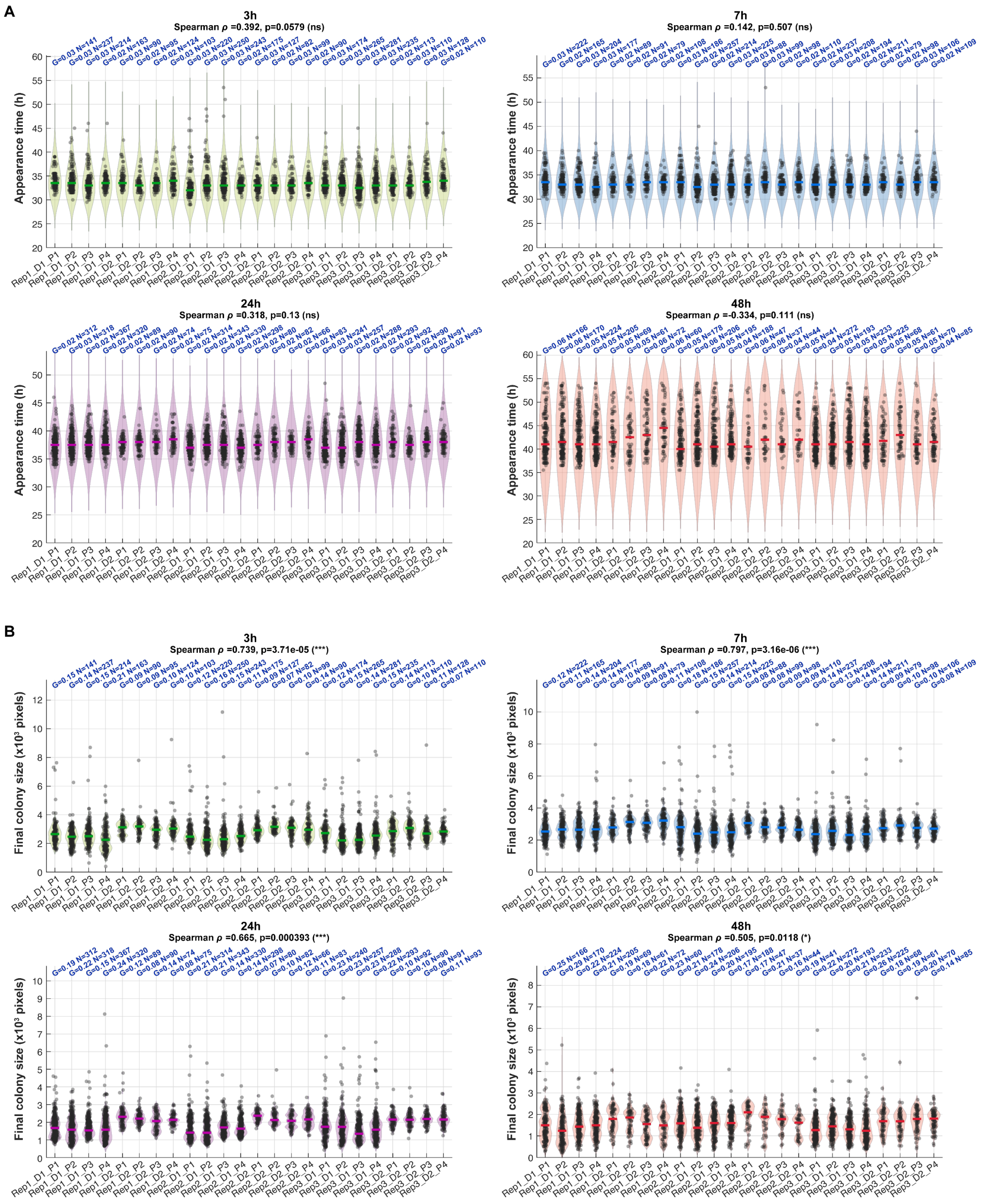


**Supplementary Figure S4. Relationship between colony number per plate and Gini coefficients for appearance time and final colony size across growth phases.** Violin plots showing **A** appearance time (h, hours) and **B** final colony size distributions for individual plates across all biological replicates (Rep), dilutions (D), and technical replicates (P) at 3h, 7h, 24h, and 48h growth phases. Gini coefficients (G) and colony counts (N) are annotated per plate in blue. Horizontal lines indicate the median per plate. Spearman $\rho$ and $p$ values reflect the correlation between colony count and Gini coefficient across all plates within each growth phase; not significant (ns), *$p < 0.05$, ***$p < 0.001$.

### Supplementary Figure S5

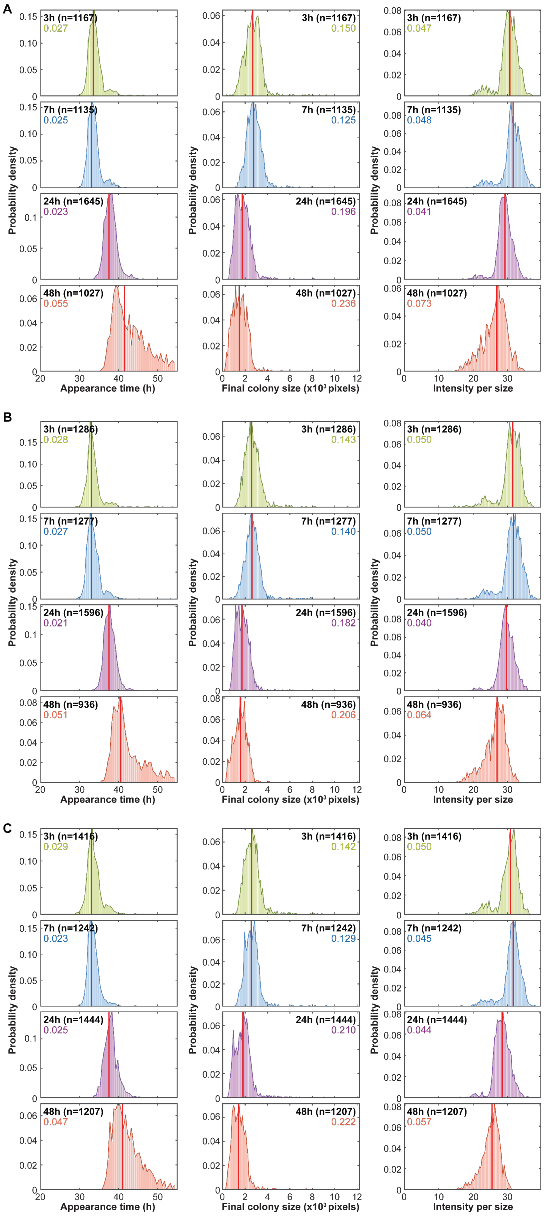


**Supplementary Figure S5. Colony phenotype distributions of *Y. pseudotuberculosis* across growth phases are reproducible between biological replicates.** Data represents the same experiment as in Fig. 2, shown for individual biological replicates. **(A-C)** Appearance time (left panels), final colony size (middle panels), and summed pixel intensity (right panels) distributions shown as probability density functions for colonies originating from cells harvested at 3, 7, 24, and 48 hours of growth. n representing the number of colonies per sample group in each individual experiment (replicate). **A–C**, Gini values are shown in color of growth condition, and median values are indicated by red vertical lines.

### Supplementary Figure S6

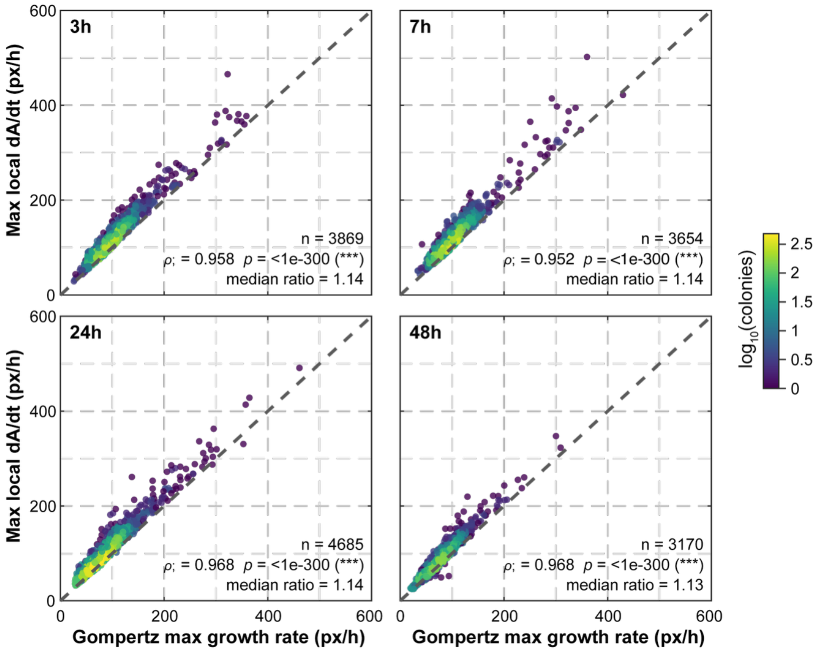


**Supplementary Figure S6. Comparison of model-free and Gompertz-derived maximum growth rates.** Each point is one colony, plotted as the maximum growth rate from the Gompertz fit (μ_max = A_max·k/e, px/h; x-axis) against the maximum local dA/dt (px/h; y-axis). The maximum local dA/dt is the maximum slope of the colony area trajectory A(t), taken as a local linear regression slope over a sliding window of five consecutive timepoints. Both estimates were computed from the same smoothed area trajectories, for the same colonies, in the in vitro sample groups (n shown per panel). Points are coloured by the $\log_{10}$ number of colonies in their local density bin (40 × 40 bin grid), and the dashed line is the identity line (y = x). Each panel reports the Spearman correlation (ρ) with its two-sided *p*-value, and the median ratio of max local dA/dt to Gompertz μ_max across colonies.

### Supplementary Figure S7

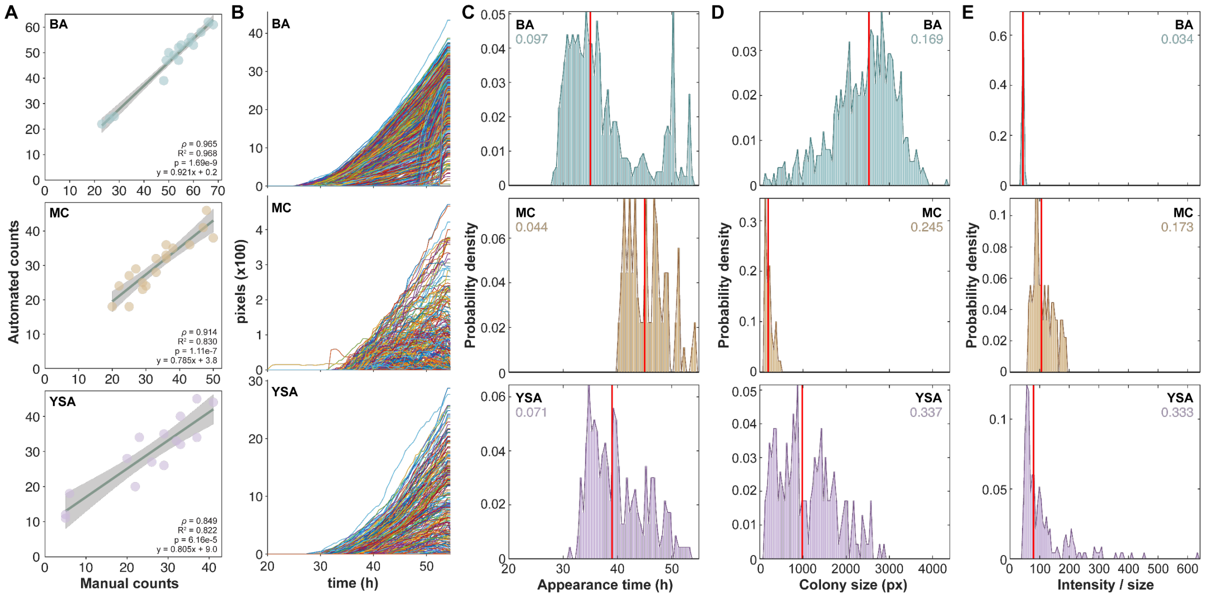


**Supplementary Figure S7. Assessment of HeteroTyper across diverse dark agar media using colony morphologies.** For dark media experiments, wt cells grown for 48-hours were inoculated into rich liquid medium at 26°C. Samples collected at 48 hours were serially diluted, and the $10^7$ dilution was spread onto 6 plates for each dark media (BA, MC, YSA), incubated at room temperature for 20 hours, and imaged in the HeteroTyper system. Each panel row corresponds to a distinct medium, with BA in the top, MC in the middle, and YSA in the bottom panel. **A** Comparison of manual and automated colony counts across dark media, demonstrating the accuracy of the computational analysis. **B** Growth curves of colonies grown across diverse dark media; each line represents an individual colony, with data combined across all replicates per media type. **C** Appearance time, **D** final colony size, and **E** summed pixel intensity per colony size, shown as probability distribution functions for colonies grown on BA (n = 752), MC (n = 545), and YSA (n = 439). For each parameter in **C–E**, Gini values are shown in color of growth condition, and median values are indicated by red vertical lines.
